# Optimized mucin-selective enrichment strategy to probe the mucinome

**DOI:** 10.1101/2023.12.18.572204

**Authors:** Keira E. Mahoney, Vincent Chang, Taryn M. Lucas, Krystyna Maruszko, Stacy A. Malaker

**Affiliations:** Department of Chemistry, Yale University, New Haven, CT 06511, USA

**Author notes:** To whom correspondence should be addressed: Stacy A. Malaker.

## Abstract

Mucin-domain glycoproteins are densely O-glycosylated and play critical roles in a host of healthy and disease-driven biological functions. Previously, we developed a mucin-selective enrichment strategy by employing a catalytically inactive mucinase (StcE) conjugated to solid support. While this method was effective, it suffered from low throughput and high sample requirements. Further, the elution step required boiling in SDS, thus necessitating an in-gel digest with trypsin. Here, we optimized our previous enrichment method to include elution conditions amenable to mucinase digestion and downstream analysis with mass spectrometry. This increased throughput and lowered sample input while maintaining mucin selectivity and enhancing glycopeptide signal. We then benchmarked this technique against different O-glycan binding moieties for their ability to enrich mucins from various cell lines and human serum. Overall, the new method outperformed our previous procedure and all other enrichment techniques tested. This allowed for effective isolation of more mucin-domain glycoproteins, resulting in a high number of O-glycopeptides, thus enhancing our ability to analyze the mucinome.

## Introduction

Mucin-domain glycoproteins are characterized by extremely dense O-glycosylation that contributes to a unique secondary structure which exerts biochemical and biophysical influence on cells.^1^ The canonical family of mucins, e.g. MUC2 and MUC16, bear massive mucin domains that can reach 5-10 MDa in size, while many other proteins contain smaller mucin domains.^2–4^ Indeed, we recently introduced the human “mucinome” which comprises nearly 350 proteins predicted to bear the dense O-glycosylation characteristic of mucin domains.^2^ For instance, lubricin is a mucin-domain glycoprotein present in synovial fluid and plays an important role in joint lubrication and synovial homeostasis.^5^ Protein tyrosine phosphatase receptor type C (PTPRC), also known as CD45, is an essential regulator of T- and B-cell receptor signaling through activation of various Src family kinases.^6^ Though these proteins are known to contain densely glycosylated regions, it is often unclear exactly how O-glycans contribute to the overall structure and function of mucin proteins. Clearly, the mucinome comprises rich biology that we are just beginning to uncover.

Our previous work indicated that a catalytically inactive point mutant of a mucin-selective protease, StcE (StcE^E447D^), retained its binding specificity for mucin domains while leaving them intact for subsequent analysis.^2,7–9^ Thus, we conjugated StcE^E447D^ to POROS-AL beads which generated a solid phase support material to use for enrichments. Samples were incubated with beads overnight at 4 °C, washed, and bound proteins were eluted by boiling in SDS. Enriched proteins were separated via SDS-PAGE and subjected to ingel digest. We used this procedure to identify mucin-domain glycoproteins from several cancer-associated cell lines and crude ovarian cancer patient ascites fluid.^2^ While effective, the method was accompanied by several key drawbacks. Given that the interaction between StcE^E447D^ and the mucin targets was strong, harsh elution conditions (i.e., boiling in SDS) were required, which necessitated an in-gel digest to remove the detergent prior to analysis via mass spectrometry (MS). Eight bands were cut per lane, and two lanes were run for each replicate, for a total of 96 gel slices per experiment. To process these samples required nearly five days at the bench, followed by nearly five days on the instrument. Further, in-gel digests are not amenable to mucinase or O-glycoprotease digestions, thus many of the mucin domains likely remained undigested and were not subjected to MS analysis. Finally, we required approximately 3 mg of cell lysate and/or ascites fluid, thus limiting the type and quantity of samples that could be processed.

Beyond StcE, many other mucinases and O-glycoproteases have been introduced to the field.^8–12^ Indeed, we recently published an “enzymatic toolkit” comprised of six mucinases from various commensal (e.g., *Akkermansia muciniphilia, Bacteroides thetaiotaomicron*) and pathogenic (e.g., *Streptococcus pneumoniae, Escherichia coli*) bacteria.^8^ More recently, we characterized an enzyme from *Serratia marcescens* called SmEnhancin (SmE) and demonstrated that it cleaves N-terminally to glycosylated Ser or Thr residues bearing a myriad of different glycan structures.^11^ Interestingly, most of these enzymes bear distinct globular domains with varying functionalities.^9,11^ Naturally, they all contain an active site domain that engages in proteolytic cleavage, but many also contain “mucin-binding domains” (MBDs) that are reportedly responsible for binding to mucin proteins. In fact, deletion of StcE’s MBD (called X409) completely abrogated StcE binding to mucin-producing cells in human tissues.^13^ SmE contains two putative MBDs with unknown specificity and function.^11^ Given the differences in cleavage motifs, structures, and MBDs between StcE and SmE, we reasoned that these could act as complementary mucin enrichment tools.

Here, we sought to improve our mucin-selective enrichment technique to enhance throughput, decrease sample input, and allow for the identification of O-glycopeptides. We first optimized the elution procedure, demonstrating that sodium deoxycholate (SDC) acts as a suitable and MS-compatible replacement for SDS. With this method in-hand, we enriched mucin-domain glycoproteins from human serum which allowed for near-complete sequence coverage of lubricin, along with many O-glycopeptides from other mucin proteins. Finally, we benchmarked the improved StcE^E447D^ enrichment against an inactive point mutant of SmE (SmE^E245D^), X409, and the lectin Jacalin and demonstrated that StcE^E447D^ enrichment was the most robust among them. Overall, we present an improved enrichment technique and complementary tools to probe the mucinome.

## Methods

### Mucin enrichment procedure

Mucinases and MBDs were expressed as previously described;^7,11,13^ Jacalin was purchased from Vector Laboratories. The pET28 plasmids for His-tagged SmEnhancin and StcE were kindly provided by the Bertozzi laboratory. StcE^E447D^, Jacalin, X409, and SmE^E245D^ were conjugated to NHS-Activated Beads (Pierce) and washed 3x with 1 mL of PBS, followed by the addition of the proteins, which were allowed to bind overnight at 4 °C. Free NHS-esters were capped by adding 100 mM Tris, pH 7.4 to the bead slurry for 20 min at 4 °C. BCA assays (Thermo-Fisher Scientific) were performed after the reactions in order to determine sufficient binding efficiency. After capping, beads were washed 3x with a high salt buffer (20 mM Tris, 500 mM NaCl) followed by a buffer without salt (20 mM Tris). For mock enrichment testing, 5 μg of recombinant CD45 (R&D Systems) was added to 50 μg of BSA. Pooled human serum (Innovative Research) was filtered through a 0.22 μm vacuum filter (Millipore) and the concentration was taken using a NanoDrop One spectrophotometer (Thermo Scientific). Approximately 50 μL of 50 mg/mL serum was brought to a final volume of 400 μL in 20 mM Tris and EDTA was added to a final concentration of 5 mM. Lysates from the original enrichment paper^2^ were kept stored at -80 °C; 500 μg of the K562 and Capan2 lysates were brought to a final volume of 500 μL in no salt buffer and EDTA was again added to a final concentration of 5 mM. The protein-conjugated solid supports (100 μL) were added to the samples and allowed to rotate at 4 °C overnight. After binding, the beads were washed 3x with the high salt buffer and 2x with the no salt buffer. Enriched mucins were eluted by the addition of 100 μL of 0.5% SDC in 20 mM Tris, followed by boiling the beads at 95 °C with agitation. The elution was repeated and pooled. In the detergent comparison experiments, mucins were eluted by following the Rapigest and ProteaseMAX manufacturer recommendations.

### Mass spectrometry sample preparation

Following elution, the enriched samples were subjected to reduction, alkylation, and proteolytic digestion. To begin, dithiothreitol (DTT) was added to a concentration of 1 mM and allowed to react at 65 °C for 20 min followed by alkylation in 1.5 mM iodoacetamide (IAA) for 15 min in the dark at RT. Subsequently, 800 ng of mucinase SmE was added and allowed to react overnight at 37 °C. At this point, the samples were split into two aliquots, one containing 90% and the other containing the remaining 10%. The former represented the “mucinase-only” digest that was not subjected to any further proteolysis. The latter was subjected to an additional digestion with 100 ng of trypsin (Promega) for 6 hours at 37 °C.

All reactions were quenched by adding 1 μL of formic acid (Thermo Scientific, 85178) and diluted to a volume of 200 μL prior to desalting. Addition of formic acid also caused SDC to precipitate out of solution, and the resulting supernatant was transferred to a new tube before desalting. Desalting was performed using 10 mg Strata-X 33 μm polymeric reversed phase SPE columns (Phenomenex, 8B-S100-AAK). Each column was activated using 500 μL of acetonitrile (ACN) (Honeywell, LC015) followed by 500 μL of 0.1% formic acid, 500 μL of 0.1% formic acid in 40% ACN, and equilibration with two additions of 500 μL of 0.1% formic acid. After equilibration, the samples were added to the column and rinsed twice with 200 μL of 0.1% formic acid. The columns were transferred to a 1.5 mL tube for elution by two additions of 150 μL of 0.1% formic acid in 40% ACN. The eluent was then dried using a vacuum concentrator (LabConco) prior to reconstitution in 8 μL of 0.1% formic acid.

### Mass spectrometry data acquisition

Samples were analyzed by online nanoflow liquid chromatography-tandem MS using an Orbitrap Eclipse Tribrid MS (Thermo Fisher Scientific) coupled to a Dionex UltiMate 3000 HPLC (Thermo Fisher Scientific). For each analysis, 6 μL was injected onto an Acclaim PepMap 100 column packed with 2 cm of 5 μm C18 material (Thermo Fisher, 164564) using 0.1% formic acid in water (solvent A). Peptides were then separated on a 15 cm PepMap RSLC EASY-Spray C18 column packed with 2 μm C18 material (Thermo Fisher, ES904) using a gradient from 0-35% solvent B (0.1% formic acid with 80% acetonitrile) in 60 min.

Full scan MS1 spectra were collected at a resolution of 60,000, an automatic gain control (AGC) target of 3e5, and a mass range from 300 to 1500 m/z. Dynamic exclusion was enabled with a repeat count of 2, repeat duration of 7 s, and exclusion duration of 7 s. Only charge states 2 to 6 were selected for fragmentation. MS2s were generated at top speed for 3 seconds. Higher-energy collisional dissociation (HCD) was performed on all selected precursor masses with the following parameters: isolation window of 2 m/z, stepped collision energy 25-30-40% normalized collision energy (nCE), Orbitrap detection (resolution of 7,500), maximum inject time of 50 ms, and a standard AGC target. An additional electron transfer dissociation (ETD) fragmentation of the same precursor was triggered if 1) 3 of 8 HexNAc or NeuAc fingerprint ions (126.055, 138.055, 144.07, 168.065, 186.076, 204.086, 274.092, and 292.103) were present at ± 0.1 m/z and greater than 5% relative intensity, 2) the precursors were present at a sufficient abundance for localization using ETD. If the precursor was between 300-850 m/z and over 3e4 intensity, an ETD scan was triggered and read out using the ion trap at a normal scan rate. For precursors between 850-1500 m/z with an intensity greater than 1e5, supplemental activation was used (EThcD) was also applied at 15% nCE and read out in the Orbitrap at 7500 resolution. Both used charge-calibrated ETD reaction times, 150 ms maximum injection time, and 200% standard injection targets.

### Mass spectrometry data analysis

Raw files were searched using Byonic against the human proteome. Mass tolerance was set to 10 ppm for MS1s and 20 ppm for MS2s. Met oxidation was set as a variable modification and carbamidomethyl Cys was set as a fixed modification. For most samples, we used the default O-glycan database containing 9 common structures, less H1N1A3 and N2 with two of any glycans allowed as a common modification. Serum files were searched using the N-glycan ‘57 most common human plasma’ database with one allowed as a rare modification. Three common modifications were allowed for each peptide. Files were searched with cleavage N-terminal to Ser and Thr, ‘specific’ cleavage, and six allowed missed cleavages. Samples treated with trypsin were searched with the same parameters, but also allowed cleavage C-terminal to Arg or Lys. Results were filtered to a score of >200 and a logprob of >2. Resultant protein identifications were characterized as mucins if the ‘Mucin Score’ from reference [2] was higher than 1.2. The mass spectrometry proteomics data have been deposited to the ProteomeXchange Consortium via the PRIDE partner repository with the dataset identifier PXD046534. For peptide-level analyses, glycopeptide identity was manually validated, and relative abundances were obtained by generating extracted ion chromatograms and determining area under the curve. For protein level abundances, we relied on the Byonic calculated “Protein Intensity” output from search results.

Reviewer account details:

Username

Password: ft86BMBO

## Results

### Optimization of enrichment technique and elution procedure

As described above, our previous mucin enrichment procedure involved an in-gel digest which is primarily conducive to trypsin enzymatic treatment (**Figure 1A**, left). Previous attempts to perform in-gel digestion with mucinases proved to be much less successful. We attribute this to the size of the mucinases (∼100 kD) versus trypsin (∼30 kD); it is likely that the mucinases are too large to be absorbed into the gel and thus cannot perform effective proteolysis. This is a key limitation in the workflow because trypsin is generally unable to digest mucin domains due to the dense O-glycosylation and the dearth of tryptic cleavage sites.^14–16^ Thus, mucin domains will likely remain undigested in the gel and will not be identified by MS. Coupled with the inherent low-throughput of in-gel digests, we sought to optimize the elution from mucinase-conjugated beads. We reasoned that this would allow for higher throughput, lower input amounts, and subsequent mucinase digestion to identify mucin O-glycopeptides (**Figure 1A**, right).

**Figure 1.**
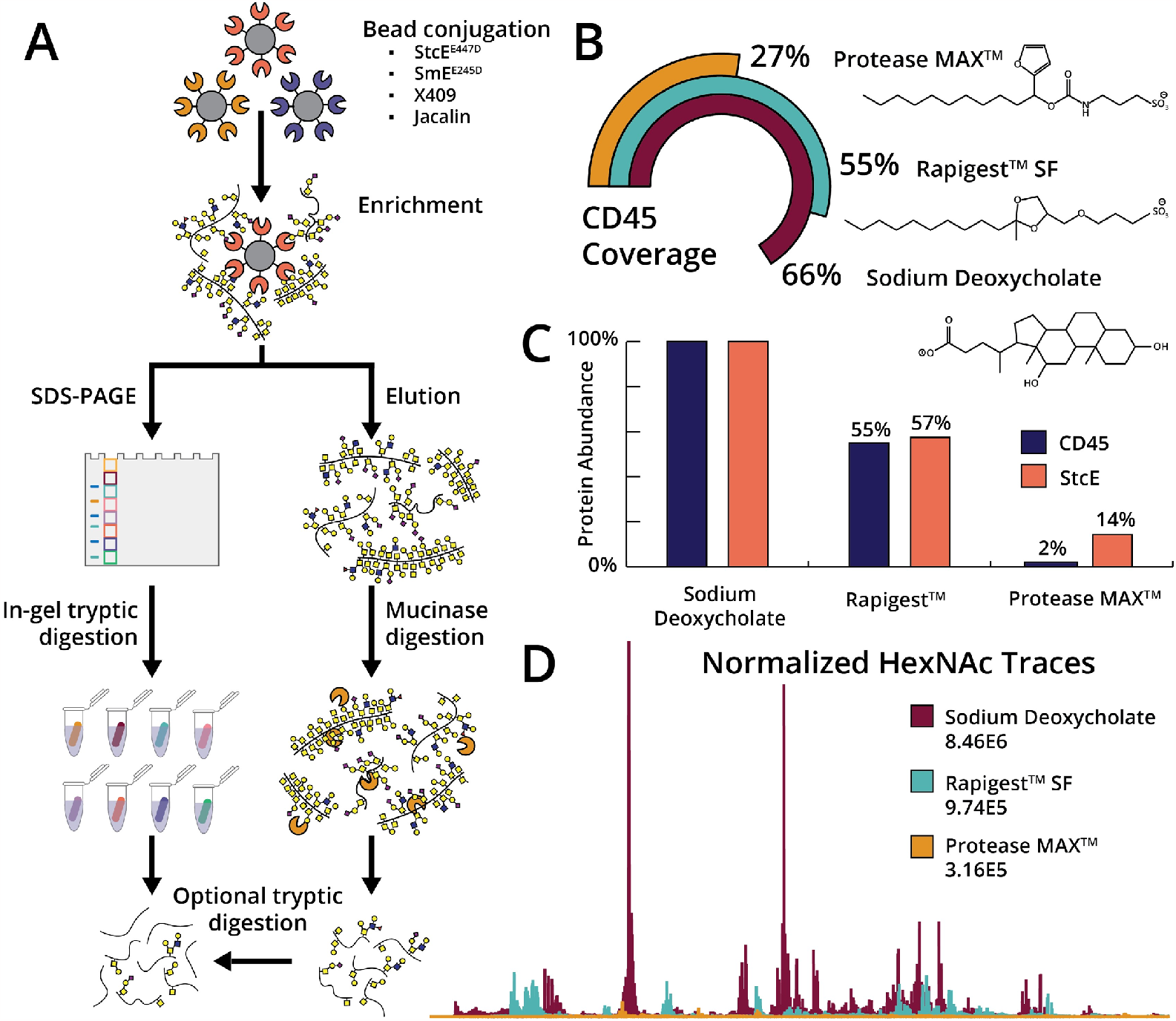
Optimization of mucin-selective enrichment techniques. (A) Mucinases, mucin-binding domains, or lectins were conjugated to NHS-ester beads and allowed to react with various complex samples. In our previous workflow (left), mucins were eluted by boiling in SDS and then subjected to in-gel digestion with trypsin. In our optimized procedure (right), we can elute with MS-friendly detergents and subject the mucins to mucinase digestion and/or tryptic digestion prior to MS analysis. (B) A mock enrichment solution containing CD45 and BSA was added to StcE^E447D^-conjugated beads and allowed to react overnight. After washing with 500 mM NaCl in 20 mM Tris three times and 20 mM Tris twice, the beads were separated into three aliquots and eluted with Rapigest, ProteaseMAX, or SDC. The elutions were subjected to digestion with trypsin and analyzed by LC-MS/MS, followed by data analysis using Byonic. Total CD45 protein coverage is depicted for Rapigest (teal), ProteaseMAX (yellow), and SDC (maroon). Inset: structures of the three detergent reagents. (C) Using protein intensities calculated by Byonic, we normalized the amount of protein to the sample with the most protein content and calculated the amount of CD45 (blue) to StcE mutant (orange). (D) Using Xcalibur software, 204.0867 extracted ion chromatograms (XICs) were generated. The same color scheme as in (B) applies here.

Prior to evaluating elution conditions, we optimized two other aspects of the procedure. First, we previously employed POROS-AL beads for conjugation, wherein beads were reacted with StcE^E447D^ via reductive amination followed by capping in Tris-HCl. While effective, this method exhibited a few drawbacks: (1) the reaction proceeds after the addition of sodium cyanoborohydride, which can be fatal if ingested, inhaled, or absorbed through the skin, and (2) their small size renders it difficult to completely remove supernatant without disturbing the bead pellet. Thus, we turned to N-hydroxysuccinimide (NHS) solid support, which allowed for coupling in mild conditions (e.g., PBS at neutral pH), followed by capping in 100 mM Tris-HCl. The NHS-conjugated beads were easier to handle and resulted in very similar results when compared to the POROS-AL beads.

Secondly, we formerly observed that StcE^E447D^-conjugated beads enriched a total of 75 proteins from HeLa lysate, 28% of which were mucin-domain glycoproteins.^2^ This proved to be more selective for mucins than Jacalin, which is a well-established lectin that enriches O-glycoproteins bearing truncated O-glycans. However, the 72% of protein identifications from non-mucins indicated room for improvement with regard to non-specific binding. Therefore, we hypothesized that more stringent wash steps prior to elution could remove more nonmucin proteins and enhance the overall selectivity. Toward this end, we reasoned that high salt conditions would interrupt electrostatic interactions that could cause nonspecific binding, as typically employed in many immunoprecipitation and protein-based enrichment procedures. After incubating mucinase-conjugated beads with various samples, we washed the beads three times in 500 mM NaCl in 20 mM Tris, followed by two washes in 20 mM Tris to remove the salt. This resulted in an improved selectivity for mucins by removing proteins that were non-specifically bound to beads.

Finally, we moved to optimize the elution conditions to circumvent the necessity for in-gel digestion. We investigated detergents Rapigest and ProteaseMAX, which are well established in the field as MS-compatible detergents that have structural homology to SDS, thus should perform similarly in maintaining protein solubility during boiling. However, we quickly realized that the costs associated with each (Rapigest ∼$66/mg, ProteaseMAX ∼$76/mg) would be prohibitive in large-scale experiments. Thus, we investigated other potential detergents and discovered SDC which is a chaotropic bile salt detergent reported to be especially useful in disrupting and dissociating protein interactions, especially with regard to elution or regeneration of affinity columns. Similar to Rapigest and ProteaseMAX, digestion efficiency is enhanced by SDC (unlike SDS), thus does not need to be removed before digestion with trypsin. This ultimately lowers the number of necessary steps and improves yield from enrichment. Further, we purchased SDC for $1 per gram (∼$25/25 g), representing 6600- and 7600-fold reductions in price when compared to Rapigest and ProteaseMAX, respectively. To compare the effectiveness of all three detergents, we generated a mock enrichment consisting of mucin-domain glycoprotein CD45 spiked into BSA. The sample was allowed to incubate with StcE^E447D^-conjugated beads overnight prior to separation into three aliquots of equal volume. Elution was performed in the respective detergents, followed by reduction, alkylation, and proteolytic digestion. As shown in **Figure 1B**, elution with SDC resulted in the highest sequence coverage of CD45 (66%) when compared to elution with Rapigest (55%) and ProteaseMAX (27%). The increased sequence coverage was concomitant with enhanced protein abundance, however, led to a higher amount of StcE leaching from the beads (**Figure 1C**). Additionally, HexNAc (i.e., GlcNAc and GalNAc) subjected to collision-based fragmentation techniques produces a characteristic HexNAc oxonium ion peak at 204.0867 m/z. We thus surveyed the HexNAc oxonium ion profiles to quickly assess success in enriching CD45 from the BSA solution. As demonstrated in **Figure 1D**, the 204 trace for ProteaseMAX is replete of signal, whereas both Rapigest and SDC resulted in a high abundance of 204 ions. That said, the overall levels of signal were highest in SDC, suggesting that it is the most effective elution buffer. We concluded that the best conditions for mucin enrichment involved a combination of NHS beads for conjugation, high salt washes, and SDC-based elution.

### Application of improved enrichment to human serum enhances mucin and O-glycopeptide identification

With these techniques in-hand, we next applied our enrichment technique to pooled human serum. Key to this analysis was a mucinase recently introduced by our laboratory, SmE, which strongly outperformed commercially available O-glycoproteases in the MS analysis of mucin-domain glycoproteins.^11^ Here, we subjected human serum to enrichment with StcE^E447D^ and the eluent was digested with SmE and trypsin followed by LC-MS/MS analysis (“StcE^E447D^ Enriched”). For comparison, we treated the serum with SmE and trypsin without any prior enrichment (“Unenriched”). The data was searched using Byonic against the human proteome and filtered for scores >200 and logProb >2. As demonstrated in **Figure 2A** (top), a large portion of the protein identifications in both samples were globulins (teal) and lipoproteins (pink). In our previous work, we developed a “Mucin Domain Candidacy Algorithm” which considers predicted O-glycosites, glycan density, and subcellular location in order to output a “Mucin Score”. This value was developed as a method to gauge the likelihood that a protein contains a mucin domain; Mucin Scores higher than 2 represented those with a high probability, between 2-1.5 were considered medium confidence, and 1.5-1.2 were low confidence.^2^ Using these values, we determined the number of protein identifications considered to be mucins in both samples. Unsurprisingly, the percentage of mucins identified in the enriched sample was much higher, increasing from 5 to 22% of total protein identifications (**Figure 2A**, bottom). Additionally, investigate the number of proteins in each sample that were predicted to be O-glycoproteins in Uniprot. Here, we measured a dramatic improvement, increasing from a total of 37 (17%) to 56 (31%) after enrichment (**Figure 2A**, bottom). We note that Uniprot is largely underannotated for O-glycosylation and thus our numbers are likely an underestimation of the true number of O-glycoproteins in the samples.

**Figure 2.**
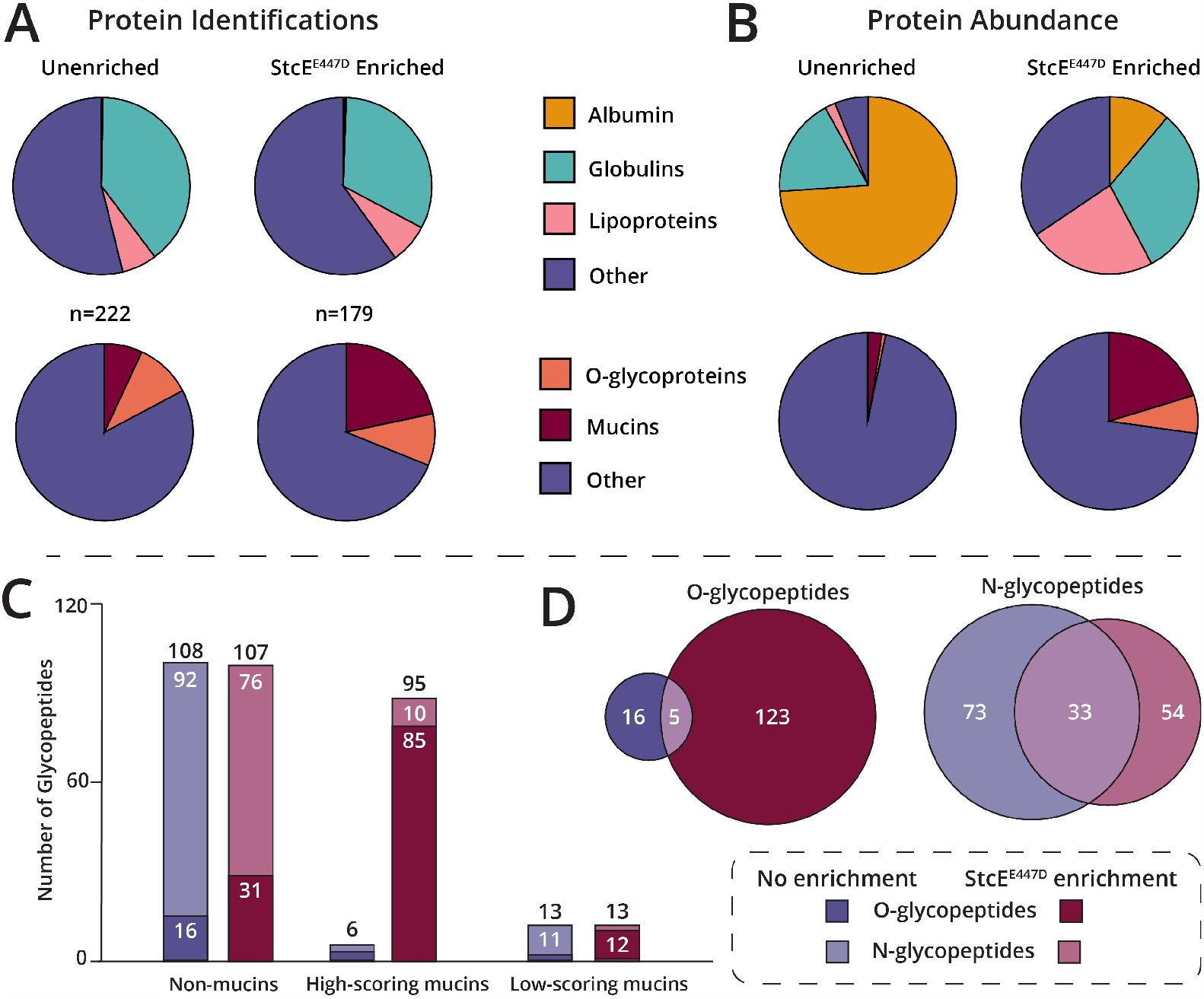
Enrichment of human serum using optimized StcE^E447D^ enrichment procedure. Human serum was digested with SmE and trypsin (“Unenriched”), and also subjected to enrichment followed by digestion with mucinase SmE and trypsin (“StcE^E447D^ Enriched”). Resultant peptides were analyzed by LC-MS/MS on a Thermo Orbitrap Eclipse Tribrid using an HCD-pd-ET(hc)D method. Raw data was searched with Byonic against the human proteome; for more detailed information, see Methods. (A) Protein identifications were analyzed by GO term classification. Top: Albumin (yellow), globulins (teal), lipoproteins (pink), and other (purple) protein distribution in unenriched and enriched samples. Bottom: Mucin scores and O-glycoprotein predictions were used to evaluate the total number of mucins (maroon), O-glycoproteins (orange), or other (purple) identified in both samples. (B) Protein abundances were calculated with Byonic “protein intensity” and labeled with the sample color scheme as in (A). (C) Glycopeptide identities were investigated using Mucin Scores where “non-mucins” were considered anything with a score of <1.2, high scoring mucins had a score >2, and low scoring mucins had a score between 1.2-2. Those glycopeptides identified without enrichment are plotted in purple, those found after StcE^E447D^ enrichment are plotted in maroon; darker shades indicate O-glycopeptides, lighter shades represent N-glycopeptides. (D) Comparison of N-and O-glycopeptides identified in enriched (maroon) and unenriched (purple).

Next, as opposed to evaluating the total list of proteins, we sought to investigate the various abundances of each protein using Byonic calculated “protein intensity”.^18^ In this analysis, nearly 75% of total protein abundance in the unenriched sample could be attributed to albumin (**Figure 2B**, top). The StcE^E447D^ enrichment caused this to drop to 11%, which likely allowed for the identification of proteins at a much lower abundance. To be sure, the total abundance of mucins combined with O-glycoproteins improved from 7.5% to over 25% (**Figure 2B**, bottom). Thus, by simultaneously depleting albumin and enriching mucins, we can dramatically improve our ability to detect these species.

We next investigated the glycoproteomic landscape in human serum pre- and post-enrichment. Here, we again took into account Mucin Scores to consider how many glycopeptides were identified from “non-mucins” (Mucin Score <1.2), “low-scoring mucins” (Mucin Score 1.2-2), and “high-scoring mucins” (Mucin Score >2; **Figure 2C**).^2^ Here, without enrichment (purple), 85% of the detected glycopeptides originated from non-mucins. After mucinselective enrichment, this number decreased to 50%, whereas the number of glycopeptides originating from high-scoring mucins improved from 5 to 44%. Additionally, the total number of glycopeptides increased nearly two-fold, from 127 to 215 after enrichment. Taking this one step further, we examined the breakdown of N-versus O-glycopeptides in the two samples (**Figure 2D**). In the unenriched serum, the grand majority of glycopeptides were modified by N-glycans (83%, 106/127). After enrichment, we observed a dramatic increase in the number of O-glycopeptides (128 of 215 total) as well as a shift in the ratio of N-to O-glycopeptides (41% N-, 59% O-glycopeptides). While all of the aforementioned experiments involved enzymatic digestion with SmE and trypsin, previously, we demonstrated that SmE-only digestion could result in deep glycoproteomic coverage of mucin domains larger than 100 amino acids. The lack of workhorse protease treatment results in fewer unmodified peptides that can contribute to ion suppression of glycopeptides, thus improving glycoproteomic sequencing.^11,16^ Thus, a portion of the serum enrichment was subjected to SmE digestion and the results are shown in **Supplemental Figure 1**. In this experiment, we demonstrated that an even higher number of the identified proteins were mucins and O-glycoproteins (72%), and the overall abundance of these proteins accounted for approximately 50% of total protein abundance. Additionally, we observed complementarity in the two digests with regard to the O-glycosites and unique glycopeptides identified (**Supplemental Figure 2**). Thus, SmE-only digestion is useful in isolating and identifying mucin-domain and O-glycoproteins. Note that all glycopeptides detected from these experiments can be found in **Supplemental Tables 1-2**. Taken together, we demonstrated robust mucin enrichment from human serum, which allowed improved detection of both mucin glycoproteins and O-glycopeptides.

### Glycoproteomic analysis of lubricin enabled by StcE^*E447D*^ enrichment and SmE digestion

Lubricin, also known as proteoglycan 4, and encoded by gene *PRG-4*, is a mucin-domain glycoprotein secreted in the synovial joint and is important for cartilage integrity.^5,19,20^ In healthy joints, lubricin molecules coat the cartilage surface, providing boundary lubrication and preventing cell and protein adhesion.^21–23^ Arthropathies such as osteoarthritis (OA) and rheumatoid arthritis (RA) are concomitant with altered lubricin glycosylation; glycomic studies have demonstrated a decrease in sialic acid and a higher prevalence of truncated O-glycan structures.^19,20^ Glycoproteomic sequencing of lubricin would provide atomic-level insight into these noted glycan aberrations. Previously, this was accomplished by subjecting synovial fluid (SF) to ion exchange chromatography, in-gel digestion, and cotton-based glycopeptide enrichment. The resultant peptides were then subjected to sialidase and O-glycanase treatment, which trimmed the O-glycan structures to mono-or disaccharides at each site. Using these procedures, a total of 168 O-glycosites were identified from 185 unique glycopeptides.^5^ While this was an impressive accomplishment, the therapeutic relevance for the procedure is limited by the need for a high amount of starting material, several time-consuming processing steps, and enzymatic glycan removal. Additionally, SF is relatively invasive to obtain; a serum-based sequencing method could be beneficial to develop diagnostic tools for OA/RA patients. However, while lubricin is present at 5-7 μg/mL in SF, it is much less abundant in serum (0.5-1.4 μg/mL), thus enrichment is necessary.^24,25^

We reasoned that our optimized StcE^E447D^ enrichment procedure could allow for the glycoproteomic mapping of lubricin from human serum. To do so, 2.5 mg of human serum was reacted with StcE^E447D^-conjugated beads and eluted with SDC. From there, 10% of the sample was treated with mucinase SmE followed by trypsin. Given that lubricin has ∼1000 amino acids in its mucin domain, we digested the remaining sample (90%) using SmE only. As demonstrated in **Figure 3A**, when the enriched serum was treated with both SmE and trypsin, 89% of the identified peptides were unmodified. On the other hand, when SmE was the only protease used for digestion, 71% of the total peptide content was identified as O-glycopeptides. An example of this is demonstrated in **Supplemental Figure 3**, whereby the lubricin glycopeptide **S**(GalNAc-Gal-NeuAc)TMPELNP in the SmE only digest was found at nearly 1e7 relative abundance with few co-eluting peptides, whereas the signal was suppressed to 1e6 in the SmE/trypsin digest. Additionally, the MS1 of the peptide in the latter sample was heavily chimeric, with at least 3-4 coeluting species present.

**Figure 3.**
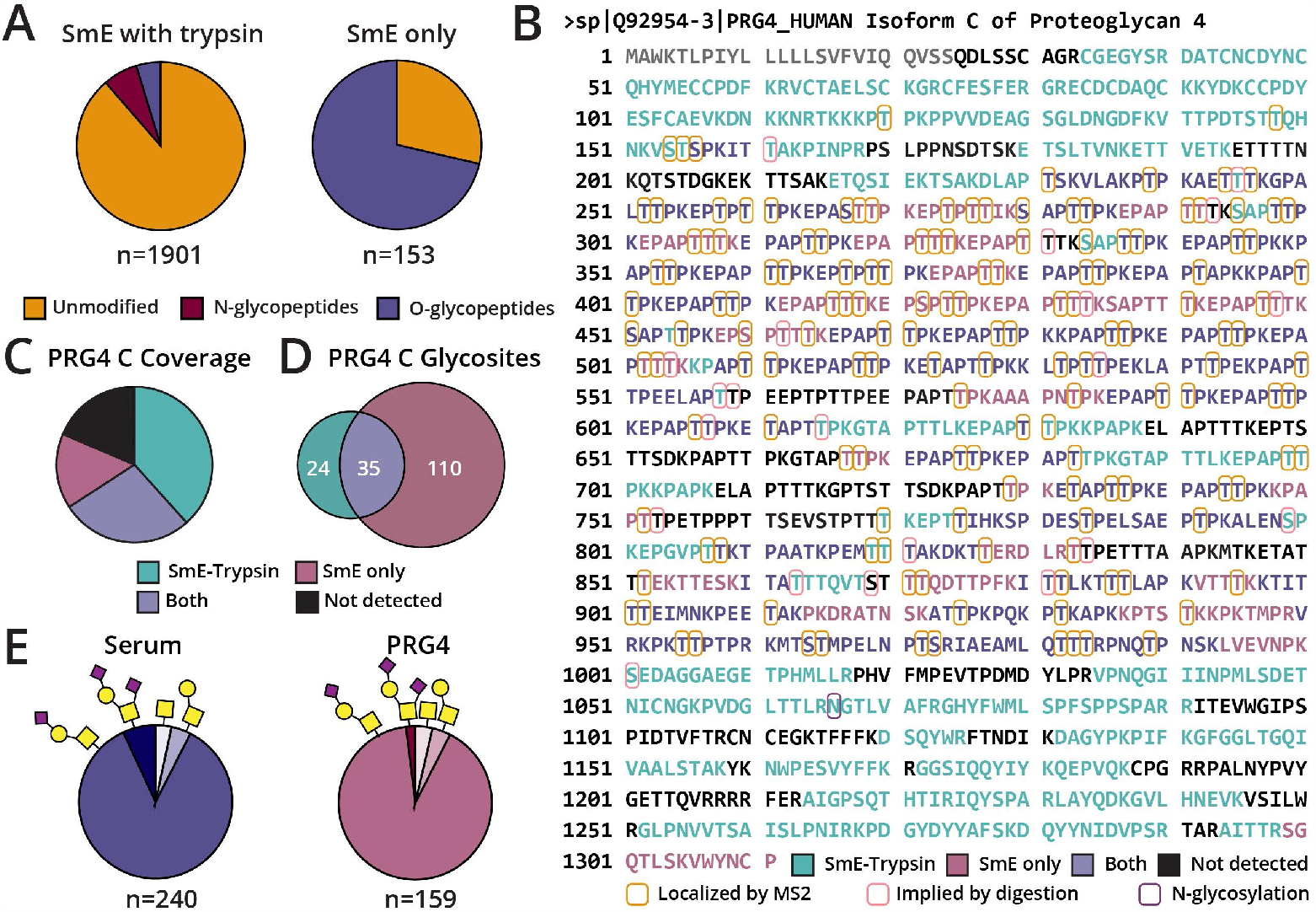
Glycoproteomic sequencing of lubricin from mucin-enriched human serum. Human serum (2.5 mg) was subjected to StcE^E447D^ enrichment, mucinase +/-trypsin digestion, and intact glycoproteomic analysis. Data was searched using Byonic against a curated mucin database and glycopeptides were manually validated. (A) Left, human serum was digested with SmE overnight and trypsin (6 h) resulting in 1901 peptides, 89% of which were unmodified. Right, human serum was digested with SmE overnight, resulting in 153 peptides identified, 71% of which were O-glycopeptides. (B) Lubricin Isoform C sequence coverage from the two digestion conditions. Peptides only identified in the SmE + trypsin digest are highlighted in teal, those from the SmE-only digest are in pink, and peptides detected in both samples are in purple. Glycosites localized by fragmentation spectra are circled in orange, those implied from cleavage with SmE are circled in pink. N-glycosites are circled in purple. (C) Sequence coverage of lubricin isoform C. (D) Total number of O-glycosites identified from lubricin isoform C. (E) O-glycans identified from total serum (left) or lubricin (right); monosialylated core 1 structures were predominant in both.

Interestingly, while the study by Li *et al*. primarily identified isoform A from SF, we only were able to detect isoform C from serum (**Figure 3B**).^5^ Isoform C differs from the canonical PRG-4 in that it lacks amino acids 107-199 present in the isoform A sequence. As shown in **Supplemental Figure 4**, we sequenced a peptide that comprises amino acids 96–108, which can only be present in isoform C due to the deletion of residues found in isoform A; this could suggest that the different isoform expression is location and/or cell-type dependent. Regardless, in total, we attained an 82% sequence coverage of lubricin (similar to the earlier study), 66% using SmE and trypsin and 43% using SmE alone (**Figure 3C**). Additionally, we identified 169 O-glycosites (**Figure 3D**), 21 of which were implied from SmE cleavage. These numbers rival the study by Li *et al*., and we note that these glycopeptides were not subjected to any glycosidase treatment, thus are more difficult to detect and site-localize. As demonstrated in **Supplemental Figure 5**, the previous work identified 28 unique glycosites when compared to our study; however, we detected 94 unique to our study. Finally, we investigated the glycans associated with the glycopeptides in serum (**Figure 3E**) and found that the grand majority of glycopeptides were modified by a monosialylated core 1 structure, supporting previous observations. The total number of glycopeptides from enriched serum is demonstrated on the left; glycopeptides identified specifically from lubricin are shown at right. Taken together, the optimized enrichment procedure can allow for near-complete glycoproteomic sequencing of lubricin from serum. Future efforts will be devoted to applying this technique to OA/RA patient serum to improve diagnostic and/or therapeutic strategies.

**Figure 4.**
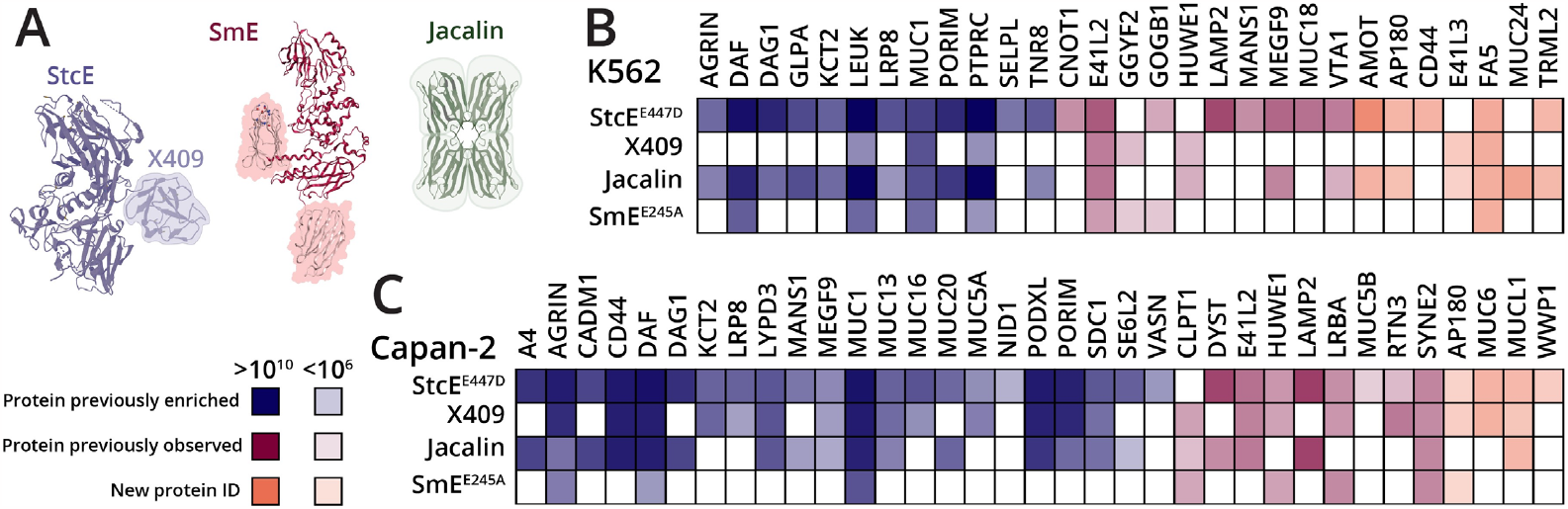
Mucin-selective enrichment from cell lysates Capan2 and K562. (A) Crystal structure of StcE (blue) and Jacalin (green), along with AlphaFold predicted structure of SmE (maroon). Accessory mucin-binding domains are indicated by blue and pink highlighting. Mucin-domain glycoproteins enriched from Capan2 (B) and K562 (C) cell lysates using StcE^E447D^, X409, Jacalin, and SmE^E245D^. Proteins were conjugated to NHS-ester beads and lysates were allowed to bind overnight at 4 °C. Enrichment elutions were subjected to SmE and trypsin digestion followed by intact glycoproteomic analysis. Data were searched using Byonic against the human proteome and/or a curated mucin database, then filtered for scores >200 and logProb >2. Abundances were calculated using “protein intensity” from Byonic output. Mucin proteins previously identified in reference 2 are depicted in blue, those that were not identified are shown in orange, and those identified but without statistical significance are shown in pink.

### Comparison of StcE^*E447D*^ enrichment to other mucin-selective isolation techniques

In the original mucinome paper, we used the inactive point mutant of StcE to enrich mucins from cancer-associated cell lines (HeLa, Capan2, K562, OVCAR-3, and SKBR3) and crude ovarian cancer patient ascites fluid. We also compared the selectivity of StcE for mucin-domain glycoproteins to the lectin Jacalin, as it has been used for decades to enrich O-glycoproteins and peptides. We showed that while Jacalin was able to enrich a higher number of mucin proteins, it also isolated a large number of non-mucins, demonstrating its broad affinity for O-glycoproteins as opposed to mucins specifically.^2^ Since then, other research has suggested that the MBD of StcE, also called X409, is able to bind mucins without the active site domain (**Figure 4A**, purple).^13^ Finally, as discussed above, we recently introduced SmE as an addition to the ever-growing mucinase toolkit; SmE cleaves in a complementary manner to StcE and likely has different endogenous targets. Further, SmE has two MBDs that appear to work cooperatively to bind mucin proteins (**Figure 4A**, maroon).^11^ We note that the insoluble nature of SmE MBDs precluded their use in this study. Taken together, we sought to compare the optimized StcE^E447D^ enrichment to X409, Jacalin, and an inactive point mutant of SmE (SmE^E245D^) to determine the complementarity and robustness of the various proteins.

Previously, we used 3 mg of cell lysate split into 6x 500 μg aliquots for StcE^E447D^ enrichment; these were then run on SDS-PAGE and two lanes of peptide elution were combined per replicate. Using this procedure, in a Capan2 cell lysate enrichment, we detected a total of 30 mucin-domain glycoproteins as statistically significantly enriched in the elution when compared to lysate alone; for K562, we enriched a total of 20 mucins.^2^ Using the improved StcE^E447D^ enrichment strategy, we analyzed 500 μg (i.e., 6x less material) of K562 and Capan2 lysate; these results are shown in **Figure 4B** and **4C**. Many of the mucins were identified in both StcE^E447D^ enrichments (22 – Capan2, 12 – K562); a heat map of these proteins’ abundances is shown in blue. In addition, we observed several “new” mucins that were not enriched previously, these proteins’ abundances are depicted in orange. In the Capan2 enrichment we identified 4 “new” mucins and in K562 we identified 7. Finally, some proteins were identified in the former experiment but were not statistically significant, these proteins are highlighted in pink. In total, we identified 35 mucin-domain glycoproteins from Capan2 and 29 from K562. Results from these experiments can be found in **Supplemental Tables 3** (Capan2) and **4** (K562). Overall, the improved enrichment technique allowed for a larger number of mucins to be identified with less sample input and much higher throughput.

Interestingly, despite claims that X409 can bind mucin glycoproteins as well as intact StcE^E447D^,^13^ we found that it was less successful at isolating mucins from these cell lines. In the Capan2 enrichment, X409 isolated a total of 23 mucin glycoproteins, many of which were present at a lower abundance when compared to StcE^E447D^; with K562, X409 only isolated a total of 8 mucins. Together, this suggests some level of cooperativity exists between the MBD and the active site domain.^26^ With Jacalin, we observed that it was again able to enrich mucin glycoproteins, but this time was not able to outperform StcE^E447D^ with respect to the number of mucins identified; this is likely attributable to the improvements in our method. Finally, we used SmE^E245D^ to enrich both cell lysates with the hope that the complementarity in cleavage motifs and accessory domains would allow for isolation of unique mucin glycoproteins. Here, we did detect a few unique mucins, but at a much lower abundance and with fewer overall protein identifications. Ultimately, StcE^E447D^ outperformed all of the other enrichment techniques in enabling the identification of mucin glycoproteins from cell lysates.

## Conclusions

Here, we present an optimized mucin-selective enrichment procedure that leverages a new elution detergent and in-solution mucinase treatment. With this improved method in-hand, we enriched human serum using StcE^E447D^-conjugated solid support and greatly increased the number of mucin glycoproteins identified as well as the number of O-glycopeptides detected. In the same experiment, we were able to obtain 80% sequence coverage and 169 unique O-glycosites from the clinically-relevant lubricin protein. Finally, we benchmarked StcE^E447D^ against X409, Jacalin, and SmE^E245D^ in enrichment of mucin proteins from cell lysates, demonstrating that StcE^E447D^ enrichment is the most robust and selective amongst them. Future efforts will be devoted to applying this technique to various patient samples to identify potential biomarkers and/or therapeutic targets.

## Acknowledgements

The authors would like to thank Jeffrey Shabanowitz for helpful discussions during the course of this work, and for members of the Bertozzi laboratory (Kayvon Pedram, D. Judy Shon) for generating the initial K562 and Capan2 cell lysates. K.E.M. and T.M.L. are supported by Endowed Fellowships in the Biological Sciences. V.C. and K.M. are supported by NSF GRFP awards. The work was supported, in part, by NIGMS R35-GM147039 and a Women’s Health Research at Yale Pilot Award.

## C.O.I. Statement

S.A.M. is a co-inventor on a Stanford patent related to the use of mucinases as research tools.

