## Supplemental Information for "Optimized mucin-selective enrichment strategy to probe the mucinome"

This file contains:

- Supplemental Figures 1-5
- Supplemental Table 2

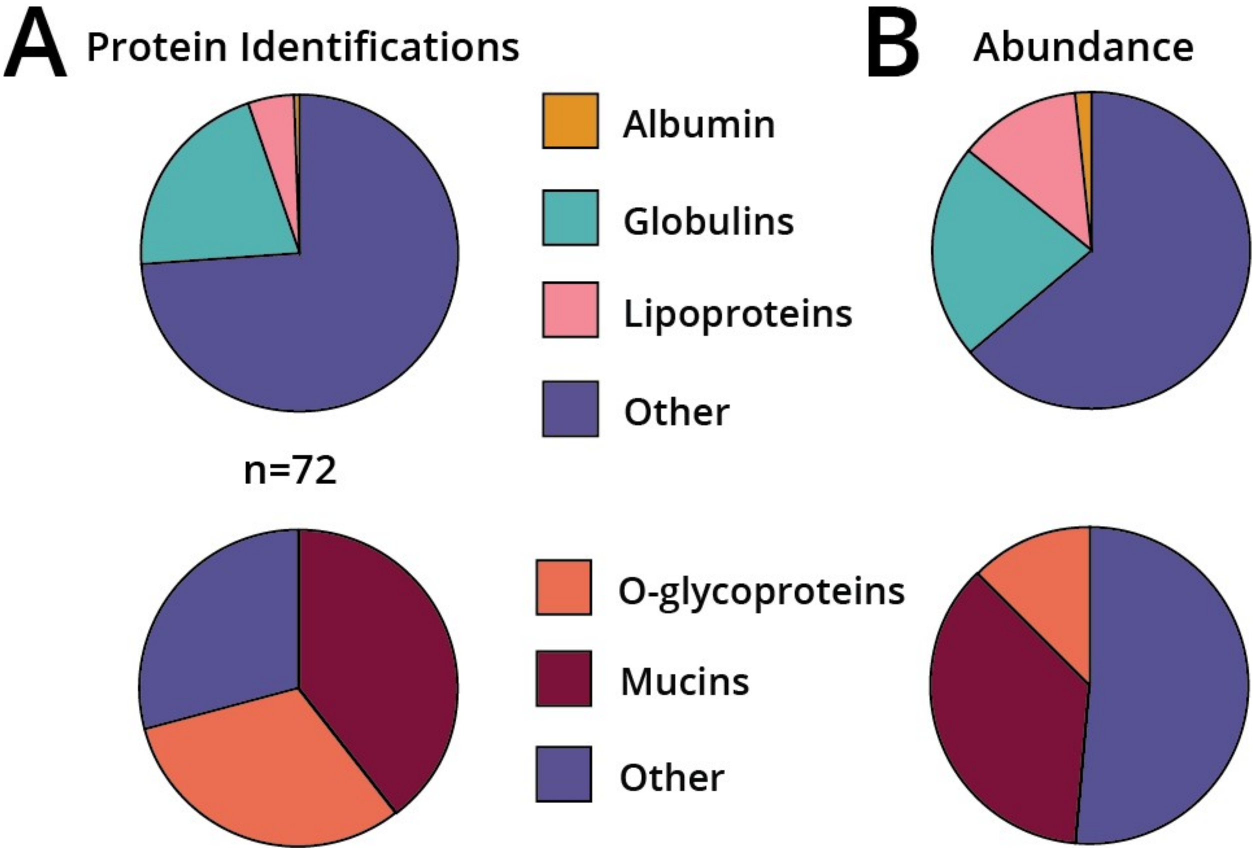


**Supplemental Figure 1. Enrichment of human serum using optimized StcE$^{E447D}$ enrichment procedure followed by digestion with mucinase SmE.** Human serum was subjected to enrichment followed by digestion with mucinase SmE. Resultant peptides were analyzed by LC-MS/MS analysis on a Thermo Orbitrap Eclipse Tribrid using an HCD-pd-ET(hc)D method. Raw data was searched with Byonic against the human proteome and resultant peptides were filtered for scores >200 and logProb >2. (A) Protein identifications were analyzed by GO term classification. Top: Albumin (yellow), globulins (teal), lipoproteins (pink), and other (purple) protein distribution in unenriched and enriched samples. Bottom: Mucin scores and Uniprot O-glycoprotein annotations were used to evaluate the total number of mucins (maroon), O-glycoproteins (orange), or other (purple) identified in both samples. (B) Protein abundances were calculated with Byonic "protein intensity". The same color scheme from (A) is used here.

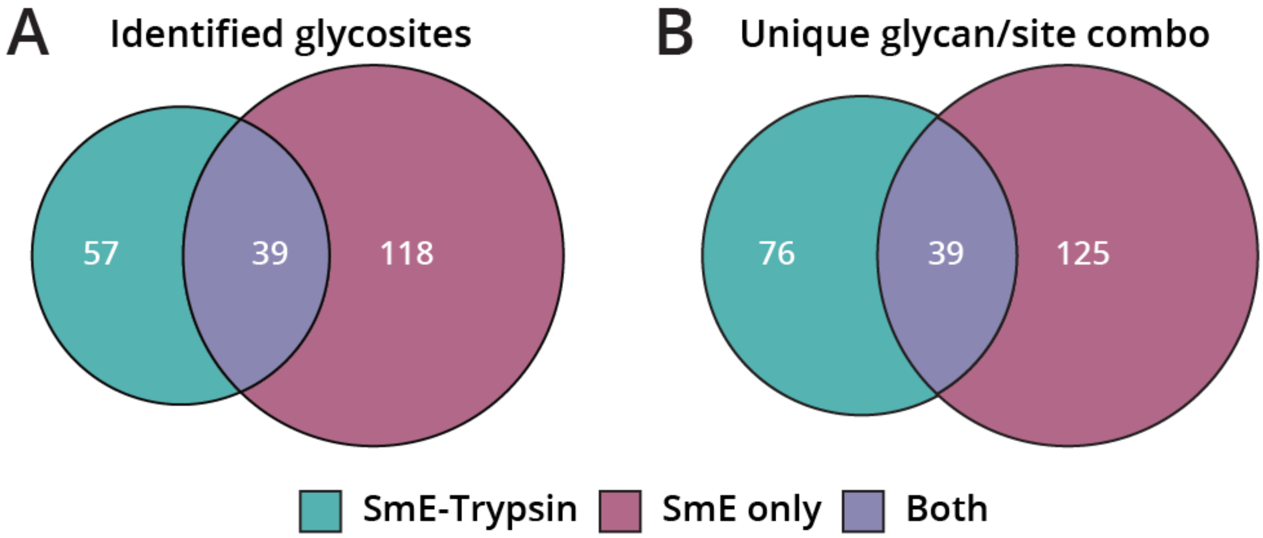


**Supplemental Figure 2. O-glycosites and unique glycopeptides identified from human serum.** Human serum was subjected to enrichment followed by digestion with mucinase SmE, with or without trypsin. Resultant peptides were analyzed by LC-MS/MS analysis on a Thermo Orbitrap Eclipse Tribrid using an HCD-pd-ET(hc)D method. Raw data was searched with Byonic against the human proteome and resultant peptides were filtered for scores >200 and logProb >2. O-glycosites (A) and Unique Glycopeptides (B) identified are depicted; those found in the digest using SmE and trypsin are in green, using SmE only are in pink, and those found in both are shown in purple.

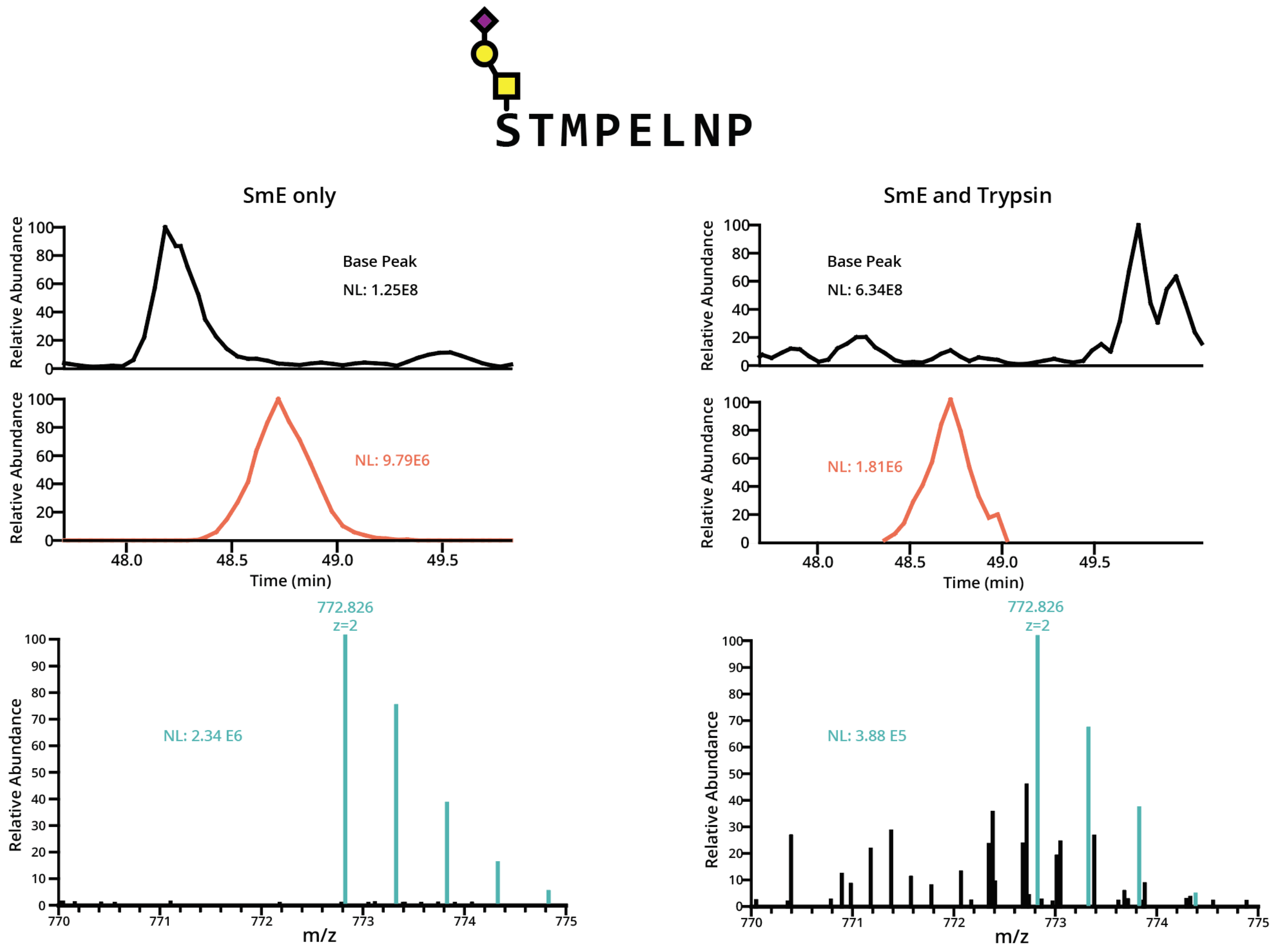


**Supplemental Figure 3. SmE-only digests improve O-glycopeptide analysis.** Pooled human serum was enriched for mucins using StcE$^{E447D}$ conjugated to NHS beads and digested with SmE only (90% of eluent, left) or SmE and trypsin (10% of eluent, right) before analysis by MS. This glycopeptide comprises amino acids 1057–1064 of lubricin and is glycosylated at Ser1057. The chromatograms and ion intensities are depicted for the base peak (black) and the extracted ion chromatogram of 772.826 (orange) corresponding to the glycopeptide mass. The MS1 of the monoisotopic precursor and its isotopes (teal) are shown along with other co-isolating species (black).

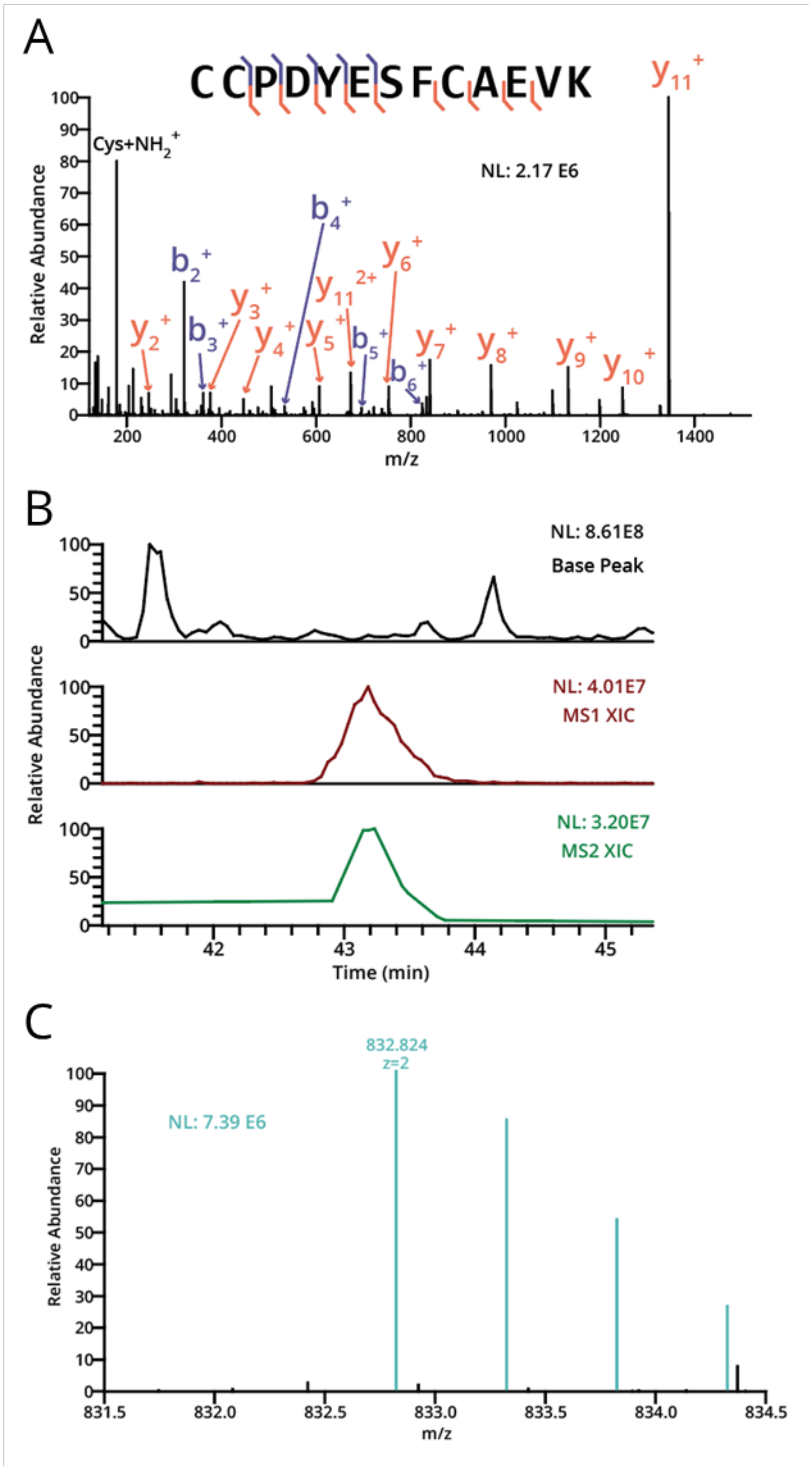


**Supplemental Figure 4. Isoform C is present in human serum. (A)** Mucin-enriched human serum was subjected to digestion with SmE and trypsin followed by MS analysis and manual data interpretation. The MS2 spectrum for the proteoglycan 4 isoform C specific peptide is shown at top. B-ions are shown in blue and y-ions in orange. (B) The chromatograms and ion intensities are depicted for the base peak (black) and the extracted ion chromatogram for the m/z of 832.824 (brown) corresponding to the peptide. A chromatogram corresponding to MS2 scans taken of the monoisotopic precursor (m/z=832.824) between 41 minutes and 45 minutes is also depicted (green). (C) The MS1 of the monoisotopic precursor and its isotopes (teal) are shown along with other weakly co-isolating species (black).

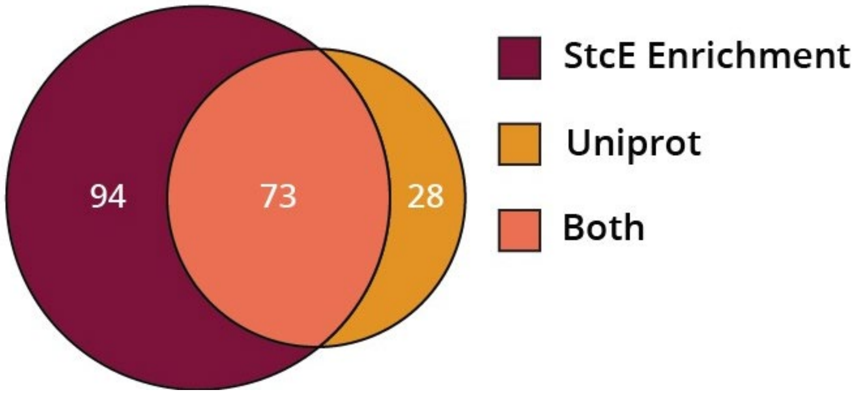


**Supplemental Figure 5. Comparison of previously reported O-glycosylation sites in lubricin and the current study.** In our study, human serum (2.5 mg) was subjected to StcE$^{E447D}$ enrichment, SmE with and without trypsin digestion, and intact glycoproteomic analysis. Data was searched using Byonic against a curated mucin database and glycopeptides were manually validated. For the glycosites reported in Uniprot, please refer to Li *et al.* for detailed methods.

**Counts**

| | No enrichment | | Enrich (SmE-tryp) | | Enrich (SmE only) | |
|---|---|---|---|---|---|---|
| | Intensity | IDs | Intensity | IDs | Intensity | IDs |
| Albumin | 8.67E+11 | 1 | 1.58E+10 | 1 | 11612807 | 1 |
| Globulins | 2.12E+11 | 93 | 4.38E+10 | 58 | 8.56E+08 | 16 |
| Lipo | 2.68E+10 | 15 | 3.33E+10 | 13 | 1.97E+08 | 9 |
| Mucin | 1.41E+10 | 11 | 2.88E+10 | 39 | 1.62E+09 | 26 |
| O-glycoproteins | 7.48E+10 | 37 | 3.87E+10 | 56 | 2.91E+09 | 35 |
| Total | 1.17E+12 | 223 | 1.41E+11 | 179 | 4.1E+09 | 72 |

**Percentages**

| | No enrichment | | Enrich (SmE-tryp) | | Enrich (SmE only) | |
|---|---|---|---|---|---|---|
| | Intensity | IDs | Intensity | IDs | Intensity | IDs |
| Albumin | 73.8% | 0.4% | 11.2% | 0.6% | 0.3% | 1.4% |
| Globulins | 18.0% | 41.7% | 31.0% | 32.4% | 20.9% | 22.2% |
| Lipo | 2.3% | 6.7% | 23.6% | 7.3% | 4.8% | 12.5% |
| Mucin | 1.2% | 4.9% | 20.3% | 21.8% | 39.7% | 36.1% |
| O-glycoproteins | 6.4% | 16.6% | 27.4% | 31.3% | 70.9% | 48.6% |

**Supplemental Table 2.** Calculated intensities and identification numbers for various protein classes. Human serum was digested with SmE and trypsin ("Unenriched"), and also subjected to enrichment followed by digestion with mucinase SmE and trypsin ("StcE$^{E447D}$ Enriched"). Resultant peptides were analyzed by LC-MS/MS analysis on a Thermo Orbitrap Eclipse Tribrid using an HCD-pd-ET(hc)D method . Raw data was searched with Byonic against the human proteome and resultant peptides were filtered for scores >200 and logProb >2; intensities were calculated from Byonic's "Protein Intensity" values. These tables were used to generate Fig 2.
